# DILS: Demographic Inferences with Linked Selection by using ABC

**DOI:** 10.1101/2020.06.15.151597

**Authors:** Christelle Fraïsse, Iva Popovic, Clément Mazoyer, Bruno Spataro, Stéphane Delmotte, Jonathan Romiguier, Étienne Loire, Alexis Simon, Nicolas Galtier, Laurent Duret, Nicolas Bierne, Xavier Vekemans, Camille Roux

## Abstract

We present DILS, a deployable statistical analysis platform for conducting demographic inferences with linked selection from population genomic data using an Approximate Bayesian Computation framework. DILS takes as input single-population or two-population datasets (multilocus fasta sequences) and performs three types of analyses in a hierarchical manner, identifying: 1) the best demographic model to study the importance of gene flow and population size change on the genetic patterns of polymorphism and divergence, 2) the best genomic model to determine whether the effective size *Ne* and migration rate *N.m* are heterogeneously distributed along the genome (implying linked selection) and 3) loci in genomic regions most associated with barriers to gene flow. Also available *via* a web interface, an objective of DILS is to facilitate collaborative research in speciation genomics. Here, we show the performance and limitations of DILS by using simulations, and finally apply the method to published data on a divergence continuum composed by 28 pairs of *Mytilus* mussel populations/species.

## INTRODUCTION

Population genomic data along with efficient computational methods are becoming increasingly available, paving the way to broad-scale application of model-based inferences for understanding signatures of evolutionary processes (Hahn 2019). Neutral processes such as divergence, gene flow and changes in population size all shape patterns of genomic variation; and so demographic models attempting to reconstruct the past history of single populations or closely-related species can also serve as null models in genome scans for selection. Considering a single species, model-based inferences are valuable for example in domesticated crops for disentangling the effect of population size changes from selection on agronomic traits (Gaut et al. 2018). Two-population models allow to tackle issues on speciation genomics, where this approach provides direct testing of distinct modes of speciation (Sousa and Hey 2013), with at the two extremes a model of allopatric speciation that occurs in complete isolation and a model where speciation is opposed by continuous gene flow. This is critical to build-up a unifying picture of the genic view of speciation by quantifying the reduction in gene exchange between lineages as a function of their molecular divergence (Roux et al. 2016; Peñalba et al. 2019); and identify *in silico* genomic regions harboring speciation genes (Roux et al. 2013; Sethuraman et al. 2019), given that their barrier effects can only be detected in the presence of ongoing gene flow (see (Ravinet et al. 2017) for a review). At a broader scale, model-based inferences can be applied to ecological communities to infer, for example, the assembly history of trophically linked species (Bunnefeld et al. 2018).

Various methods have been proposed to extract such information from population genomic data. Site frequency spectrum (SFS)-based methods compute or approximate the likelihood of the allele frequency distribution from a demographic model using either the diffusion approximation (Gutenkunst et al. 2009), the moment closure (Jouganous et al. 2017) or the coalescent (Excoffier et al. 2013). While these methods are fast and can accommodate complex demographic histories, they ignore linkage information which is informative about past demography (Terhorst and Song 2015). Therefore, other methods rely on the block-wise SFS, *i.e.* the SFS of short non-recombining blocks of sequences that are unlinked to each other (Lohse et al. 2011). That way the genealogical information contained within each block is combined along the genome. Other multilocus methods can explicitly account for recombination along chromosomes therefore capturing longer range linkage disequilibrium (e.g. based on the Sequentially Markov Coalescent: Pairwise SMC, (Li and Durbin 2011); Multiple SMC, (Schiffels and Durbin 2014); SMC++ scaling to large genomic data, (Terhorst et al. 2017)); however they are still restricted to simple demographic histories excluding migration (but see MSMC-IM, (Wang et al. 2020) and diCal2, (Steinrücken et al. 2019)). Still, the flexibility of simulation-based approximate Bayesian computation (ABC) enables including recombination within unlinked blocks in multilocus inference of complex (and hopefully more realistic) evolutionary scenarios (Beaumont et al. 2002). Although more computationally expensive, the analysis of thousands of loci results in high-precision parameter estimation for most demographic scenarios (Robinson et al. 2014; Smith and Flaxman 2020).

In this paper, we present an ABC framework (DILS) building upon and extending current statistical machinery (Pudlo et al. 2015; Roux et al. 2016). Our method is flexible both in terms of the evolutionary scenarios that can be accommodated (allowing changes in population size over time, linked selection and implementing various models of migration), and type of data (multilocus fasta sequences produced e.g. from RNAseq, DNA capture or whole genome sequencing); but it also makes important assumptions such as free recombination between (blocks of) sequences, and it is restricted to a single size-change event in the past. A major improvement compared to most existing methods is decoupling the effect of linked selection and neutral history by relaxing the assumption that all loci share the same demography (Sethuraman et al. 2019; Sousa et al. 2013). We model variation in the rate of drift among loci to account for linked selection effects due to background selection (i.e. purifying selection) in low-recombination and gene-dense regions. And by explicitly modelling variation in migration rates among loci in two-population models, we can capture the effect of selection against migrants at neutral markers linked to species barriers, and so analyse further these candidate genomic regions for reproductive isolation (Roux et al. 2013).

DILS is deployable on every machine, but also offers an online platform for configuring demographic inferences based on genomic data of thousands of loci, performing them and visualizing the returned output. These advances are made possible by progress in simulator performance (Hudson 2002), reduction in the number of simulations required to train prediction algorithms (Pudlo et al. 2015) and development of computer clusters and tools facilitating parallelism (Köster and Rahmann 2012). DILS thus contributes to the nascent and promising applications of machine learning to population genomic inferences (see (Schrider and Kern 2018) for a review). Following other user-friendly ABC programs, DILS also aims to ease the use of high-performance tools for non-experts in methodology (Cornuet et al. 2008; Cornuet et al. 2014). Importantly, as there is a limit to how much information can be extracted from genomic data, DILS also implements rigorous quality controls. Therefore, not only does the user receive 1) the best-supported model among those proposed (figure 1), 2) an estimate of the demographic parameters describing this model and 3) a locus-specific test to identify barriers to gene flow (when relevant); the user will also get feedback on whether the best model is relevant and to which extent the estimates are able to reproduce the observed data.

**Fig. 1.**
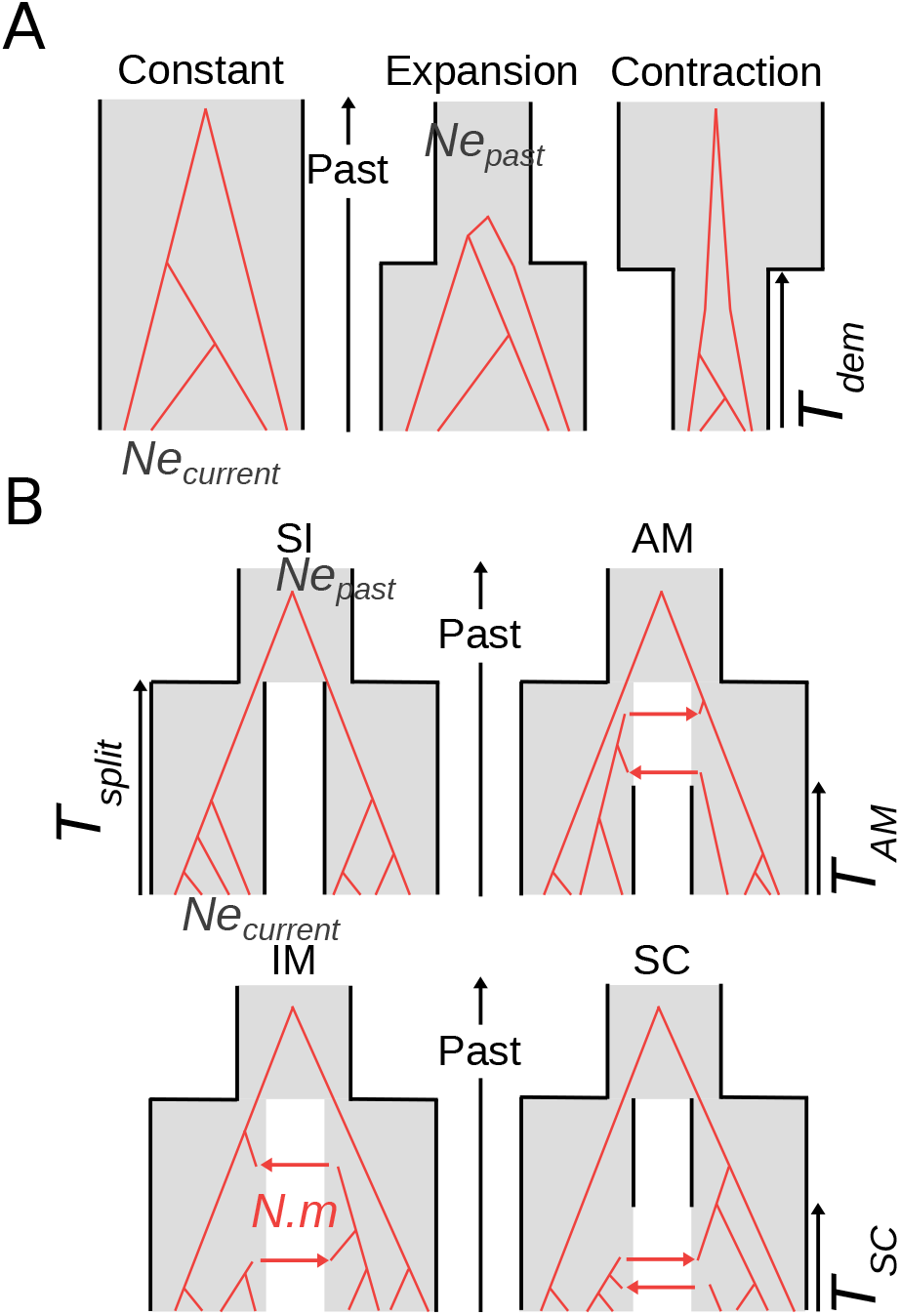
Demographic models currently implemented in DILS. **A.** Single-population models. Demographic changes occurring *T*_dem_ generations ago are modeled backwards in time by sudden transitions from *Ne*_current_ to *Ne*_past_, either for expansions or contractions. **B.** Two-population models. The Strict Isolation (SI) and Ancient Migration (AM) models are characterized by an absence of ongoing migration. Conversely, the Isolation with Migration (IM) and Secondary Contact (SC) models describe two populations that are currently connected by introgression events at rate *N.m*. The two-population models shown here are of constant size, but DILS optionally incorporates alternative versions of the same four models where effective size can change independently in both daughter populations between the present time and *T*_split_. This is a relevant addition given the influence of over-time size-changes on demographic inferences in speciation scenarios (Momigliano et al. 2020).

**Fig. 2.**
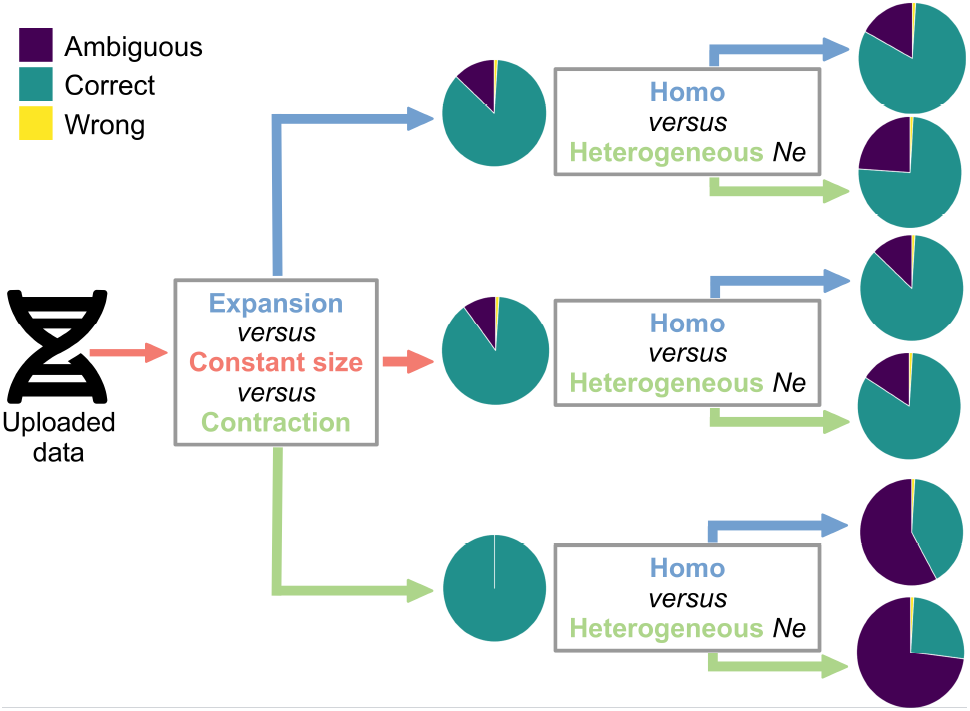
DILS performance for hierarchical comparison of single-population models. DILS first performs a comparison of the three demographic models (Expansion *versus* Constant *versus* Contraction). In a second step, it compares two genomic sub-models (homogeneous *versus* heterogeneous genomic distribution of *Ne*) for the best supported demographic model. The pie charts designate for each model the proportion of simulations performed under the corresponding model that is strongly and correctly captured (correct: blue), strongly and incorrectly captured (wrong: yellow) and without strong statistical support for any of the studied models (ambiguous: purple). The performance of DILS was based on 10, 000 pseudo-observed datasets for each of the Expansion / Constant / Contraction demographic models. Each of these 10,000 simulated datasets are evenly distributed between the two genomic sub-models, homo and hetero *Ne*. The parameters used for the simulated datasets are randomly drawn from uniform laws, with *Ne* in [1-1,000,000] individuals and *T*_dem_ in [1-2,000,000] generations. Each simulated dataset consists of 100 loci of length 1, 000 nucleotides.

A long-term aim of DILS is to facilitate collaborative research in speciation genomics. The degree of reproductive isolation appears to follow a quasi-shared molecular clock among animals, depending on the level of net genomic divergence between lineages (Roux et al. 2016). However, for the same level of divergence, two opposite situations coexist in the so-called grey zone of speciation with, on the one hand, semi-isolated pairs capable of exchanging genetic material and, on the other hand, pairs of species that are fully reproductively isolated. Many hypotheses have been advanced to explain such a reproductive barrier contrast within the same range of molecular divergence, including differences related to life history traits (internal versus external fertilization), ecology (marine versus terrestrial organisms), reproductive systems (e.g. in plants: self-incompatibility versus self-fertilization), genome size and recombination landscape, functional redundancies in genomes, etc... Speciation is such a multi-factorial process that it seems impossible for a single research group to study these different components.

Consequently, our aim is to include in DILS a collaborative science option, allowing to feed in real time the relationship between molecular divergence and genetic isolation between lineages. Available as a choice, the sharing of the inferences made in DILS, associated with the expertise that users have about their biological model, will contribute to a long-term collaborative study aiming to better understand the speciation process. This objective is illustrated here with the analysis of 28 new pairs of mussel populations whose transcriptomes were recently published, revealing ongoing gene flow for levels of divergence greater than 2%.

In this study, we have four objectives:

1. providing a flexible and powerful demographic inference method with linked selection to analyse genome-scaled dataset in single and two-population models.
2. presenting a user-friendly tool that implements this approach and paves the way for collaborative science.
3. testing the performance and limitations of the method by using simulations.
4. applying it to an empirical dataset of *Mytilus* mussels.

## MATERIALS AND METHODS

### DILS input

DILS can accommodate any type of multilocus sequence data in fasta format. It requires a single fasta file containing all sequences obtained from all populations/species (formatted as in reads2snp, (Tsagkogeorga et al. 2012)). Missing data are encoded by ‘N’ and gaps (insertion/deletions) by ‘-’. The user can define a maximum proportion of ‘Ns’/gaps (*max_N_tolerated*) and a minimum number of treatable sites (*Lmin*) beyond which a sequence is removed from the analysis. If the number of sequences per locus and per population/species is lower than a number defined by the user (*nMin*), then the locus is discarded from the analysis. Otherwise, DILS performs a draw of *nMin* sequences (without replacement) from the set of available sequences.

Sequences (genes, DNA fragments) can be produced from a variety of sequencing techniques, the most appropriate of which are RNAseq, DNA capture and whole genome sequencing. For saving computing time with large datasets (whole genomes), it is recommended to first partition the data to generate shorter blocks of sequences (i.e. produce block-wise data of a few Kb long). Note that because DILS assumes free recombination between loci, it is important for the (blocks of) sequences to be unlinked (this can be verified by using a reference genome). In the current version, there are no haplotype-based statistics being used in the inference; therefore the SNPs do not need to be phased (i.e. the association of alleles across distinct heterozygous positions in a sequence can be arbitrary). Haploid data may also be used.

### ABC implemented in DILS

#### Summary statistics

Since ABC is a category of inferential method based on the comparison between statistics summarizing simulated and observed datasets, we first describe here the statistics computed in our framework.

We assume that users are interested in carrying out inferences from multilocus datasets. For single population models, DILS calculates for each locus: 1) the pairwise nucleotide diversity (*π*) (Tajima 1983); 2) Watterson’s *θ* (Watterson 1975) and 3) Tajima’s *D* (Tajima 1989). In addition to these three statistics, the site-frequency spectrum (SFS; (Braverman et al. 1995)) is also used to summarize the data, *i.e.*, the number of single-nucleotide polymorphism (SNP) where the derived allele is present in [2*, …, nMin*] copies in the studied population/species, where *nMin* represents the number of copies sampled for a given locus. Note that singletons are excluded from the SFS to reduce inferential artifacts related to sequencing errors. If the SFS is folded by the absence of an outgroup, then the SFS will be described by the number of SNPs where the minor allele is present in [2*, …, nMin*] copies. Overall, multilocus inferences are based on 6 multilocus summary statistics which are the means and standard deviations of *π*, *θ* and Tajima’s *D*, to which we add [*nMin* − 1] individual statistics corresponding to the sum over all loci of each bin in the SFS ([(*nMin* − 1)/2] if no outgroup is available).

For models with two populations/species, *π*, *θ* and Tajima’s *D* statistics are also calculated for each of the two samples. These are supplemented with statistics approximating the joint SFS (jSFS; (Wakeley and Hey 1997)): 1) the fraction of sites showing a fixed difference between the populations/species (*S f*), 2) the fraction of sites showing an exclusive polymorphism to a given population/species (*Sx*_A_ and *Sx*_B_) and 3) the fraction of sites with a polymorphism shared between the population/species (*Ss*). Statistics describing the divergence between the two populations/species are also calculated, including the raw (*D*_xy_; (Nei and Li 1979)) and the net (*D*_a_; (Nei and Li 1979)) divergence between the population/species, and their relative genetic differentiation measured by *F*_ST_ (Wright 1943). The use of the jSFS (without singletons) as a vector of summary statistics is an option that the user can switch on or off, to avoid cases where the jSFS is composed of a large number of bins. If the jSFS is unfolded, then this vector has a length of [*nMin*^2^ - 4] available statistics (minus 4 to remove the two bins corresponding to singletons and the two bins corresponding to the fixation of the derived or ancestral allele in both samples), and a length of 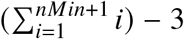 if the jSFS is folded (minus 3 to remove singletons and the (0,0) bin). The jSFS will be in the form of a square where the same number of *nMin* samples will be assumed for both species. If for a given locus the number of samples exceeds *nMin*, then a draw without replacement of *nMin* copies will be performed.

#### Simulations

In the current version of DILS, all simulations are performed using the msnsam coalescent simulator (Ross-Ibarra et al. 2008), which is a modified version of the ms program (Hudson 2002) allowing variations in sample size among loci under an infinite site mutation model. Each simulated multilocus dataset takes properties from the observed datasets (same number of genes, lengths and sample size). Since the summary statistics used to perform the ABC inferences are averages and standard deviations measured over all the surveyed loci, then for model comparisons and parameter estimations DILS randomly sub-sample 1, 000 loci (in the default version) if more loci are present in the total dataset. The purpose of this sub-sampling is to avoid unnecessarily long simulation times because the values of statistics for a given locus will not impact the used summary statistics over 1, 000 loci.

If no outgroup is provided then the same mutation rate, *μ*_*i*_ (specified by the user), is applied to all loci. Otherwise, the outgroup will be used for each locus to correct its mutation rate *μ*_*i*_ to 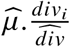 where 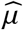 is the neutral mutation rate assumed by the user, *div*_*i*_ is the raw divergence between the focal population/species and the outgroup measured by DILS at a given locus *i*, and 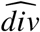 is the average raw divergence between the focal population/species and the outgroup measured over all loci. The other implication of using an outgroup will be to orientate the mutations and consequently to unfold the jSFS. Finally, a 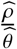 ratio value has to be specified by the user where 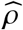 is the average population recombination rate 4*.Ne.r* (*r* in number of recombination events per generation and per nucleotide for each locus). Loci are assumed to be genetically independent from each other (unlinked), with intralocus recombination rate *ρ*.

#### Modeling linked selection

We considered that the effects of selection on linked sites can be described with a genomic model of: 1) heterogeneous effective population size (hetero-*Ne*), which well approximates the effect of background selection with a down-scaling of *Ne* (Charlesworth et al. 1993); and/or 2) heterogeneous migration rate (hetero-*N.m*) for the effect of selection against migrants (Barton and Bengtsson 1986).

In the hetero-*Ne* genomic model, the variable effective size among loci is assumed to follow a re-scaled *Beta* distribution. In other words, all populations share the same *Beta* distribution (with two shape parameters drawn from uniform distributions in [1-20]), but are independently re-scaled by different *Ne* values (drawn from uniform distributions in [0-500,000]). In the homo-*Ne* genomic model, all loci are simulated by sharing the same effective population size, which is independently estimated in all populations. This homogeneous model implies that the genome is unaffected (or equally affected) by background selection.

The hetero-*N.m* genomic model accounts for the existence of local barriers to gene flow that affects the effective migration at linked loci. We modeled variation in migration rates among loci in two ways, implemented in DILS as 2 options : 1) with a *Beta* distribution (two shape parameters drawn from uniform distributions in [1-20]), which is independently re-scaled by different *N.m* values (drawn from uniform distributions in [0-20]) in each direction of introgression; 2) with a *Bimodal* distribution where a proportion of loci (drawn from an uniform distribution in [1-100]) is linked to barriers, *i.e. N.m*=0, and the rest evolves neutrally at a rate *N.m* (drawn from an uniform distribution in [0-20]). In the homo-*N.m*, a single migration rate *N.m* (for a given direction of introgression) is shared by all loci, such that they all share the same probability to receive alleles from the other population/species.

#### Model comparisons

Here, when used alone, the term model means a given combination between a demographic and a genomic model. All comparisons are performed by using the *abcrf* function of the eponymous R package (Pudlo et al. 2015). The comparison is a two-step process.

First, grow the random forest with the *abcrf* function. This requires one reference table per model for the training. The reference table of each model is produced by 10, 000 multilocus simulations whose parameters correspond to random combinations sampled from priors. They are composed of one row per multilocus simulation and one column for each summary statistic described in section “Summary statistics”. When categories of models are compared following the hierarchical approaches (figures 2 and 3), the reference tables of the different models in the same category are merged together. For instance, in the comparison between Current isolation and Ongoing migration (figure 3), 60, 000 multilocus simulations are used for the training of the super-model Current isolation, and 80, 000 multilocus simulations for the training of Ongoing migration. Each forest is made up of 1, 000 grown decision trees regardless of the comparison made throughout the hierarchical approach.

**Fig. 3.**
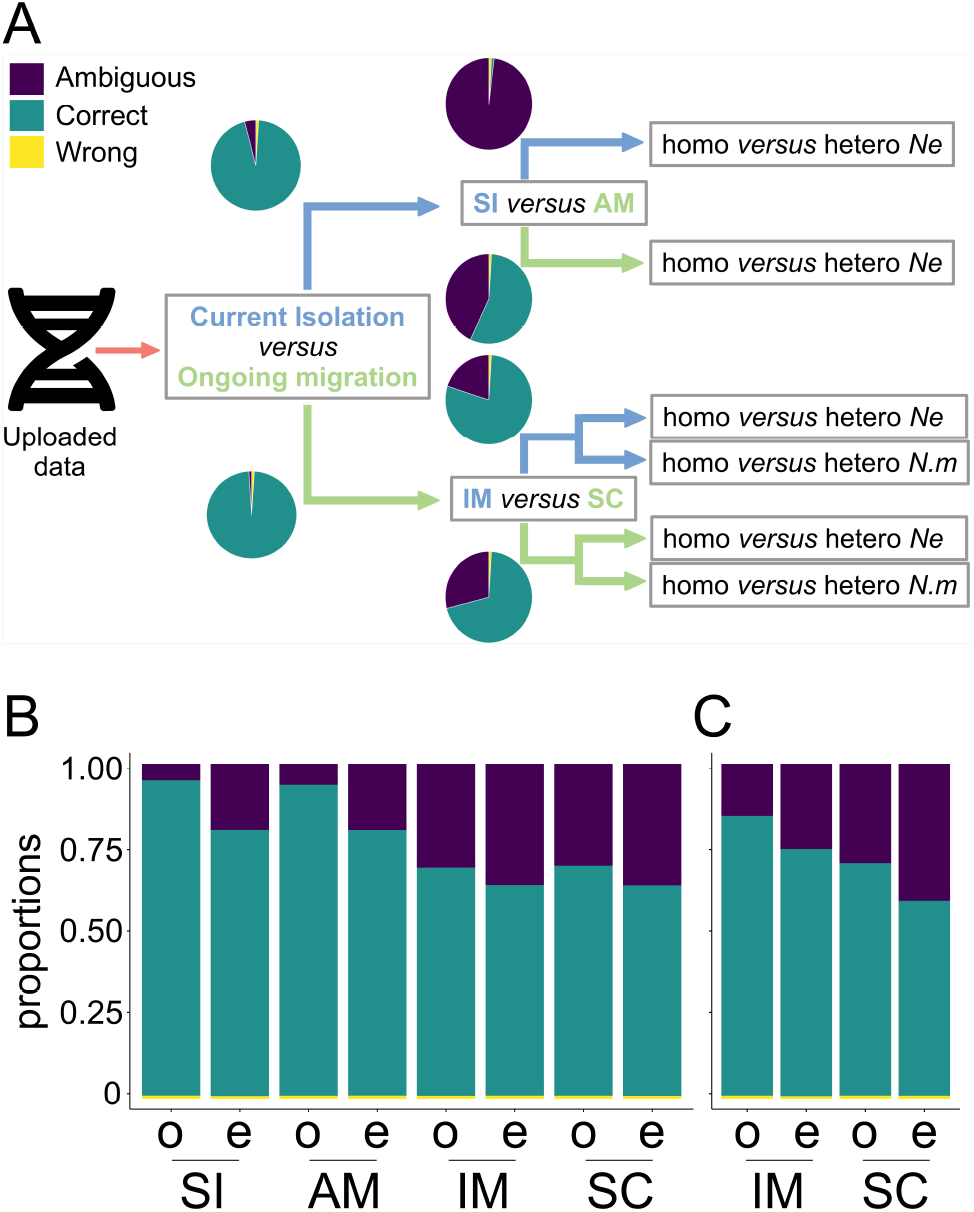
DILS performance for hierarchical comparison of two-population models. Two-population analyses are performed in three steps (panel **A**). DILS first performs a comparison of the two supermodels (“Current isolation” versus “Ongoing migration”). In a second step, it compares two demographic models for the best supported supermodel: if ‘current isolation’, then DILS tests SI versus AM; if ‘ongoing migration’, then DILS tests IM versus SC. In a third step, DILS compares genomic sub-models for variation of effective population size, *Ne* (panel **B**) and migration rate, *N.m* (panel **C**; only for ongoing migration models) thus testing for linked selection. The letters ‘o’ and ‘e’ in panels B and C indicate simulations performed under genomic h**o**mogeneity and h**e**terogeneity models, respectively. The pie charts designate for each model the proportion of simulations performed under the corresponding model that is strongly and correctly captured (correct: blue), strongly and incorrectly captured (wrong: yellow) and without strong statistical support for any of the studied models (ambiguous: purple). The performance of DILS was based on 10, 000 pseudo-observed datasets for each of the SI / AM / IM / SC demographic models. Each of these 10,000 simulated datasets are evenly distributed between the four genomic sub-models, homo and hetero *Ne* or *N.m*. The parameters used for the simulated datasets are randomly drawn from uniform laws, with *Ne* in [1-1,000,000] individuals, *N.m* in [0-20] 4*.N.m* units where *m* is the fraction of each population made up of new migrants each generation, *T*_split_ in [1-2,000,000] generations and *T*_dem_ / *T*_SC_ / *T*_AM_ between *T*_split_min_ and the sampled *T*_split_ value. Each simulated dataset consists of 100 loci of length 1, 000 nucleotides.

The second step is the prediction of the best model among those proposed by passing the observed data through the trained random forest. DILS reports the model supported by the largest number of decision trees in the random forest and its associated posterior probability.

#### Parameter estimations

Two strategies are applied to estimate the parameters describing the best-supported model among those compared. The first one is a joint estimation of the set of parameters using a rejection/regression method (Csilléry et al. 2012). Estimation is based on the 5, 000 multilocus simulations producing the statistics closest to the observed dataset among 1, 000, 000 simulations. We then correct for imperfect matches between observed and retained values of statistics. The parameter values of the selected simulations are weighted by their Euclidean distance and corrected according to a non-linear regression method using a neural network. 10 trained neural networks with 10 hidden networks are used in the regression. The second strategy is an individual estimation of each parameter by constructing a random forest of 1, 000 trees per parameter (Raynal et al. 2019).

The results from both approaches are returned to the user, as there is no evidence to further support a method over the other in terms of estimation accuracy. Within the framework of the models currently compared in DILS, both approaches produce similar estimates when tested on pseudo-observed datasets. However, joint parameter estimation has the advantages of including parameter co-variations as well as providing a probability density. This is achieved at a computational cost that is ≈ 100 times greater regarding the number of multilocus simulations, since 10, 000 are required for a parameter estimation using random forest versus 1, 000, 000 when using the rejection/regression algorithm.

#### Locus-specific model comparison

To identify barriers to gene flow among a set of sequenced DNA fragments (genes for instance), we adopt the same procedure as in *Ciona intestinalis* (Roux et al. 2013) and *Mytilus* (Roux et al. 2014), but by replacing the neural network with a random forest to divide the computational cost by 100. This step is performed by DILS only if 1) observed data better fits models with ongoing migration (IM or SC; figure 1) and 2) models of *N.m*. variation explain the data better than homogeneous models.

We first estimate the parameters of the best model from the multilocus dataset. Based on this estimation, two models are compared at each locus: 1) local-migration: the multilocus estimated model with the non-zero migration rate estimated over the whole genome; 2) local-isolation: the multilocus estimated model with a migration rate set to zero. A random forest of 1, 000 trees is then trained to recognize combinations of summary statistics specific to each of the two evaluated models. This forest allows to return for each sequenced DNA fragment the locus-specific model that best explains the statistics observed, and its posterior probability.

### DILS performance

In this study we distinguish two types of simulations. Simulations carried out to build reference tables used to train a random forest, or from which a small proportion will be sub-sampled in a rejection/regression algorithm based on the Euclidean distance with the observed data. Then a second type of simulations produces pseudo-observed datasets. These are not used for training, but to evaluate the inferential power of the ABC approach, and test whether it can recapture the parameters used to simulate the pseudo-observed datasets. To assess the reliability of model comparisons and parameter estimations, for single and two population models, we simulate pseudo-observed datasets consisting of 100 loci, of length equal to 1, 000 nucleotides, sampled from 10 diploid individuals in each population/species and a mutation rate of 5.10^−8^ mutations/nucleotide/generation. These datasets are simulated according to demographic histories using random combinations of parameters from the priors.

### Analysis with DILS of the *Mytilus* dataset

We downloaded the raw RNA-seq data deposited to the NCBI sequence read archive (BioProject ID: PRJNA560413; https://cutt.ly/OtQN1Y0) by (Popovic et al. 2019). The raw data consist in a total of ≈ 145*Gb* from the sequenced transcriptomes of 47 mussel individuals. Three individuals from the *M. californianus* species were removed as they do not belong to the *M. edulis* complex. The reference transcriptome used for the mapping is made up of 16, 151 CDS, for a total length of ≈ 23*Mb*. The reference was indexed using bowtie2 (version 2.2.6 (Langmead and Salzberg 2012)). For each individual, reads were aligned to the reference with bowtie2, and cleaned using samtools with a mapping quality threshold of 20 (version 1.3.1; (Li et al. 2009)). Individual genotypes were called using reads2snp (Tsagkogeorga et al. 2012) at positions covered by at least 8 reads, and the program outputted multilocus fasta sequences. These were then used as input data for DILS, which was run for each of the 28 possible pairs of localities by tolerating up to 20% of missing data, rejecting genes with less than 100 codons without missing data, and by keeping 6 copies per gene within each population/species. Simulations were conducted by exploring uniform priors for effective population sizes between 0 and 500, 000 diploid individuals, times of different demographic events (split, secondary contact, arrest of migration) between 0 and 1, 750, 000 generations, and migration rates between 1 and 40 4.N.m units. Presentation of the results was carried out with R (Wickham et al. 2019; Chang et al. 2019; Sievert 2018; R Core Team 2020).

## RESULTS

### Demographic and genomic models implemented in DILS

In the current version of DILS, evolutionary scenarios can be investigated for sampling schemes involving one or two populations. For both types of analysis, the first step is to compare the demographic models described in figure 1. With a single population, DILS will examine the changes in size over time. With two populations, such variations in population size are also implemented, but DILS will additionally compare alternative temporal patterns of introgression.

An important feature of DILS is to include linked selection, either through the effect of background selection that modulates the effective population size along the genome, or through the effect of selection against migrants that reduces locally the effective introgression rate in genomic regions locked to gene flow. Therefore, all demographic models exist under two alternative genomic sub-models regarding the effective population size (homo-*Ne versus* hetero-*Ne*), and the introgression rate (homo-*N.m versus* hetero-*N.m*) for models allowing migration, depending on whether these parameters are homogeneous or heterogeneous among loci.

#### Single population models

Three demographic models are compared, describing either 1) a Constant population size *Ne*_current_, 2) a recent demographic Expansion or 3) a Contraction. Demographic changes are assumed to be instantaneous, with a transition from *Ne*_past_ to *Ne*_current_ occurring *T*_dem_ generations ago (figure 1-A). The first step of the algorithm 1 implemented in DILS is to estimate the best-fitting demographic model among those depicted in figure 1-A by carrying out 10, 000 simulations under each alternative genomic sub-model (i.e. 10, 000 under the homogeneous *Ne* model, and 10, 000 under the heterogeneous *Ne* model). These simulations produce reference tables, *i.e.*, a set of simulated summary statistics used to train a random forest algorithm to predict which of the proposed models best explains observed data. The second step is to evaluate the best-fitting genomic sub-model, *i.e.* with genomic variation in effective size (hetero) or without variation (homo), for the best demographic model supported in the previous step (algorithm 1).

**Algorithm 1:**
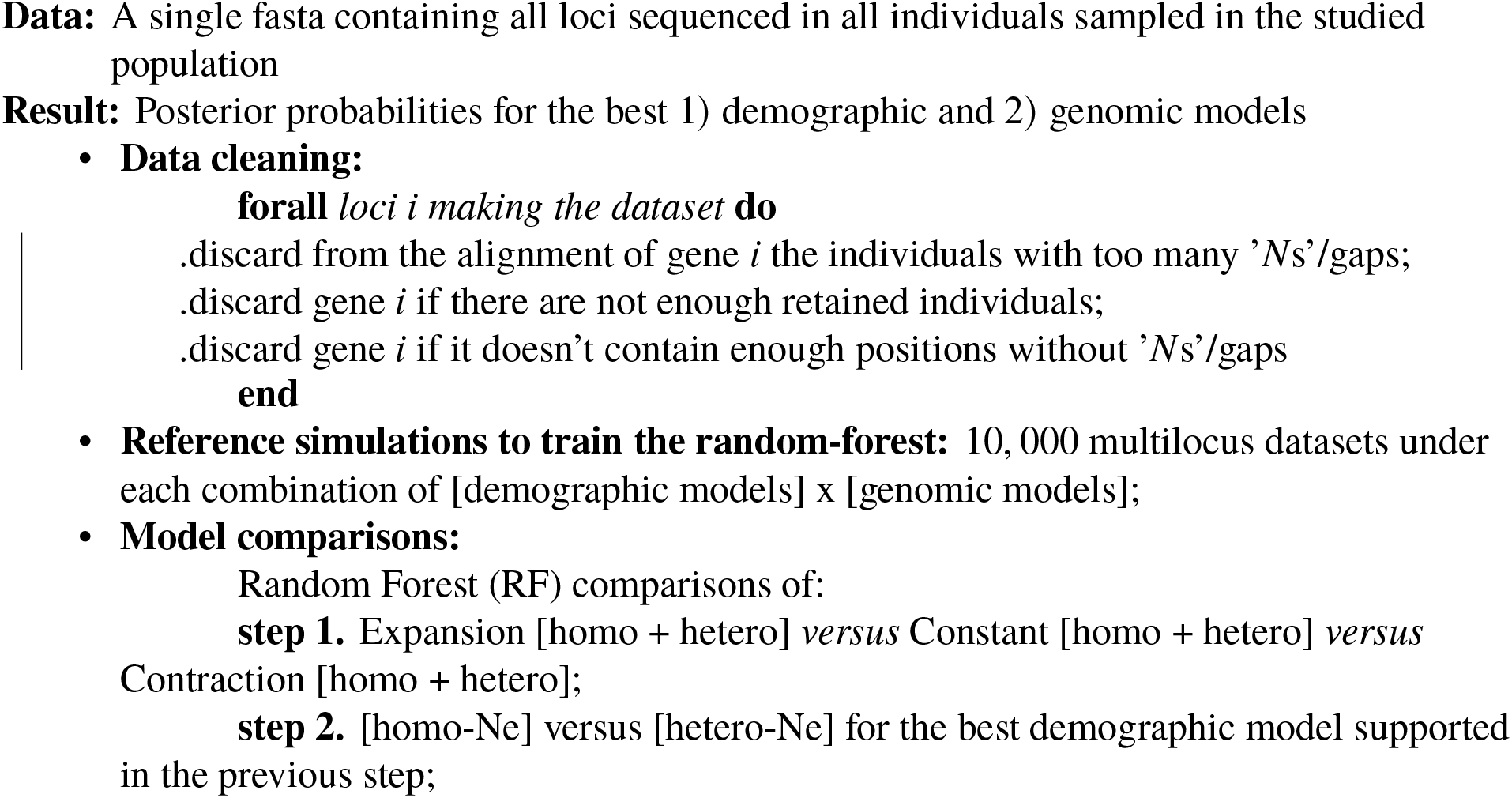
Single-population hierarchical model comparisons

#### Two population models

The two-population models are grouped into two supermodels (figure 1): with current isolation (Strict Isolation (SI) + Ancient Migration (AM)) and ongoing migration (Isolation Migration (IM) + Secondary Contact (SC)). The first step of the hierarchical comparisons therefore aims to determine which supermodel best explains the data observed in the two sampled populations (see algorithm 2). This is achieved by labeling as “isolation” all reference simulations performed under the two SI models (with homo-*Ne* or hetero-*Ne*) and the four AM models (homo-*Ne* or hetero-*Ne*, and homo-*N.m* or hetero-*N.m*). All other models are labeled as “migration” supermodel. The second step is to evaluate the demographic models described in figure 1-B (i.e. SI *versus* AM *versus* IM *versus* SC), within the supermodel that was best supported in the previous step. The third and final step is to discriminate among alternative models of genomic distribution for *Ne* (all models) and *N.m* (in case of ongoing migration).

**Algorithm 2:**
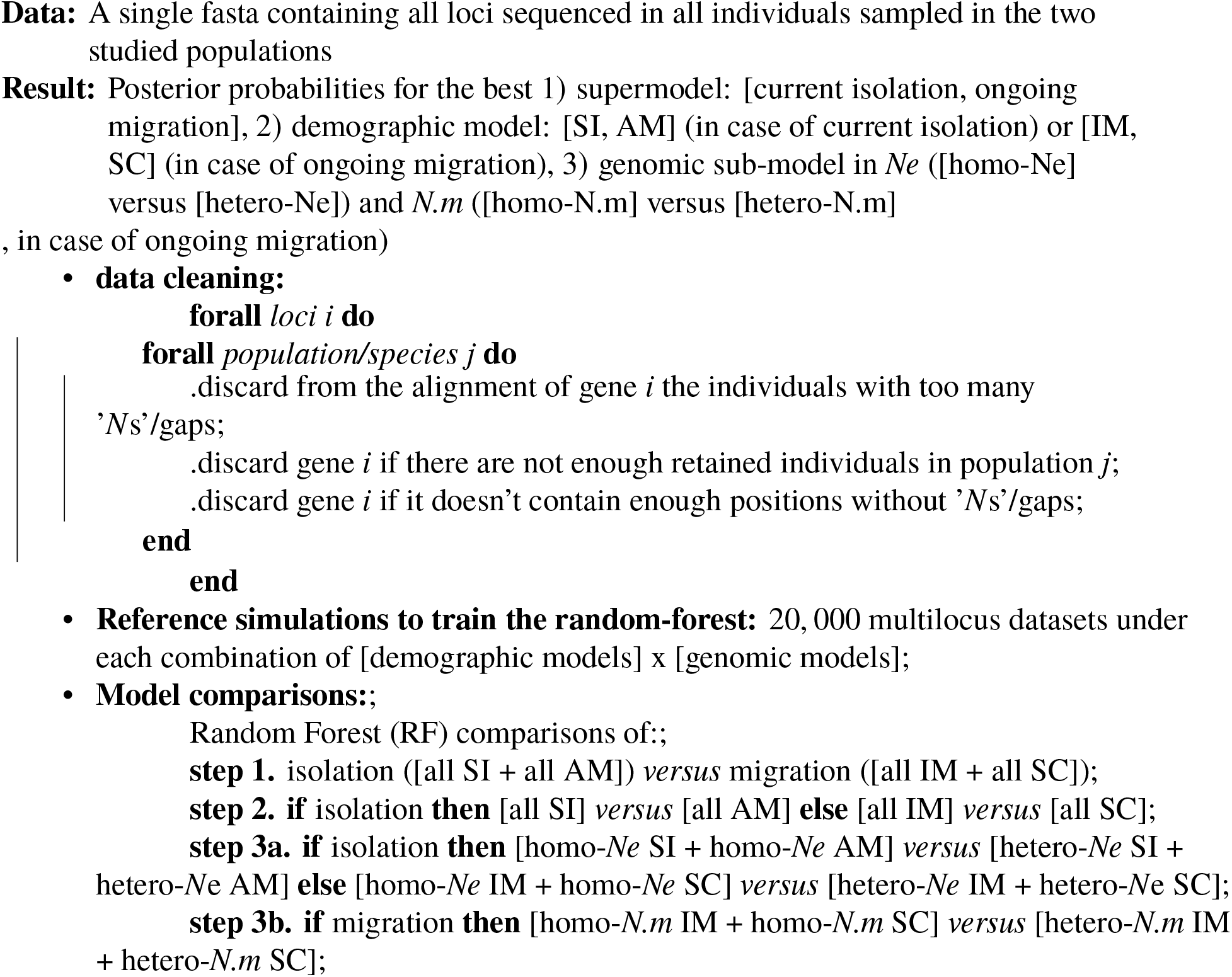
Two-population hierarchical model comparisons

### Performance for the model comparisons

In this section, we present DILS performance to compare demo-genomic models involving one (figure 2) or two (figure 3) populations/species. These evaluations were performed by analyzing pseudo-observed datasets simulated under specified models, in order to assess the efficacy of our approach to correctly support the true model. In both analyses, a given demographic model corresponds to the set of all its genomic sub-models. All model comparisons are performed using random forest algorithms (Pudlo et al. 2015; Fraimout et al. 2017), and pseudo-observed datasets do not contribute to their training.

#### Single population models

We simulated 30, 000 pseudo-observed datasets of 100 loci by using random combinations of parameters. Such a number of loci has been chosen to be conservative at a time when more than 1, 000 loci are commonly used. These simulated datasets are equally distributed between the demographic models (10, 000 for each of the three Expansion/Constant/Contraction models) and between their genomic sub-models (5, 000 for each of the two homo/hetero *Ne* alternative models). Then, for each of these pseudo-observed datasets, we apply step 1 and step 2 of the algorithm 1 in order to obtain for each model *M*, the proportion of pseudo-observed datasets that is either correctly and strongly captured by the random forest approach (blue, figure 2), falsely and strongly captured (yellow, figure 2) or ambiguous, i.e, associated to an insufficiently high posterior probability (purple, figure 2).

We determine whether an inference is strong or ambiguous from the posterior probability with which the best model is supported. For this we apply a probability threshold beyond which an inference is considered strong. This threshold is determined recursively on the basis of the false positive rate which decreases monotonically by increasing the value of the threshold. From datasets randomly simulated under different models, we establish a threshold value such that the false positive rate (*i.e.* the proportion of wrong inferences) is less than or equal to 1%. With this approach, the false-positive rate remains consistently low, but the relative proportions of true positives *versus* ambiguous cases vary according to the power of the ABC to discriminate among an arbitrary set of models.

Among the 30, 000 pseudo-observed datasets simulated under the Expansion, Constant and Contraction models, if a threshold is applied that keeps the error rate below 1%, the proportions that are correctly supported by our approach are 86%, 89% and 99% respectively, while the proportions that are ambiguous are 13%, 10% and 0.2% (figure 2). For the Expansion and Constant demographic models, the correct recapture rates of homo and hetero *Ne* genomic models range from 75% (Expansion + hetero *Ne*; 24% of ambiguity) to 86% (Constant + homo *Ne*; 13% of ambiguity). Finally, while recovering the Contraction demographic model is a very robust analysis with 99% of inferences that are both correct and associated with a high posterior probability, it is more complicated to distinguish between “Contraction + homo *Ne*” and “Contraction + hetero *Ne*”. About 41% of the pseudo-observed datasets simulated in the “Contraction + homo *Ne*” model are correctly captured by the random forest, and only 26% for the “Contraction + hetero *Ne*” model. The occurrence of a recent bottleneck tends to reduce the genomic variance of *Ne* to levels that generate apparent homogeneity (figure 2).

#### Two population models

We evaluated the performance of DILS for 60, 000 pseudo-observed datasets simulated under the “isolation” supermodel (10, 000 for each combination of [SI; AM], [homo-*Ne*; hetero-*Ne*] and [homo-*N.m*; hetero-*N.m*] for the AM model only) and 80, 000 under the “migration” supermodel (10, 000 for each combination of [IM; SC], [homo-*Ne*; hetero-*Ne*] and [homo-*N.m*; hetero-*N.m*]). As shown in figure 3, 95% of the datasets simulated under the supermodel “isolation” with random combinations of parameters from large priors are correctly recaptured by the random forest approach with a high probability (4% ambiguity and 1% error if we apply a posterior probability threshold of 0.84; table S1). Similarly, 98% of the pseudo-observed datasets under the “migration” supermodel are strongly recaptured (with 1% of ambiguity and 1% of error for a threshold of 0.665). Models with migration are globally more efficiently recaptured by DILS, relying on a lower threshold probability to be robustly supported.

The results of the performance analyses for the demographic models within each supermodel first show that a pseudo-observed dataset simulated under an SI model (homo-*Ne* and hetero-*Ne*) is very unlikely to be strongly supported in an SI *versus* AM comparison. Out of 20, 000 simulations, only 1% are correctly recaptured by DILS (98% ambiguity and 1% error for a threshold of 0.845). The AM model is more robustly supported than SI (56%), but the 10, 000 inferences made under each of the AM models lead to weak support (43% ambiguity, 1% error for a threshold of 0.705). On the contrary, the two models making the “migration” supermodel (IM and SC) are more efficiently distinguished by DILS. The 40, 000 pseudo-observed datasets randomly simulated under the IM model are captured at 79% with a high probability in the IM *versus* SC comparison (20% ambiguity, 1% error for a threshold of 0.885). Similarly, 70% of the 40, 000 pseudo-observed datasets from the SC model are correctly recaptured by DILS (29% ambiguity, 1% error for a threshold of 0.915). We then evaluate DILS performance for discriminating among alternative models of genomic distribution for the *Ne* (figure 3-B; table S2) and *N.m* (figure 3-C). Concerning the effective population size, DILS systematically recaptures the homogeneous model more easily than the heterogeneous model for each of the four demographic models tested. The most complicated model to recapture is the genomic heterogeneity of *Ne* in an SC model (≈ 64% true positives), while homo-*Ne* under an SI model is the most straightforward. Concerning the migration rate, the main parameter determining the quality of inference is the relative duration of gene flow versus speciation time *T*_split_ (figure 1). This leads to a higher robustness for the IM model (probabilities of correctly supporting homo-*N.m* and hetero-*N.m* of ≈ 86% and ≈ 76%, respectively) compared to the SC model (≈ 71% and ≈ 60%).

### Performance for the parameter estimations

In this section, we describe performance tests for estimating the parameters of different demographic models. The same procedure was applied for single and two-population models: first, simulating pseudo-observed datasets (10, 000 for the three single-population models, 2, 000 for the 14 two-population models) and then ABC estimation of the parameters to test DILS ability to recapture the parameter values used. We only detail here the results obtained for the demographic parameters, *i.e.*, those describing the mean effective population size (*Ne*_current_ and *Ne*_past_), time of split (*T*_split_), the date for the cessation of gene flow (*T*_AM_), the age of the secondary contact (*T*_SC_) and the migration rates (*N.m*).

#### Single-population models

The current effective population size is by far the parameter that is most accurately recaptured, especially in a constant and homogeneous model with a mean-squared error (MSE) close to zero (MSE≈ 0.005; figure 5-A; table S2). The introduction of a recent demographic change reduces the quality of inferences for *Ne*_current_, more for the expansion model than for the contraction model.

The inference of ancestral size conducted for 4 x 10, 000 pseudo-observed datasets shows globally a low error rate on the raw values of *Ne*_past_ with a *MSE*_max_ ≈ 0.09 (table S2). Errors depend very closely on the relative values between *Ne*_past_ and *Ne*_current_ as shown on figure 4-B with the estimates of the ratio *Ne*_past_/*Ne*_current_. Hence, while the estimate of a *Ne*_past_ is reliable for a change in size by a factor of 10, it becomes less accurate as the contrast with *Ne*_current_ is increasingly sharp.

**Fig. 4.**
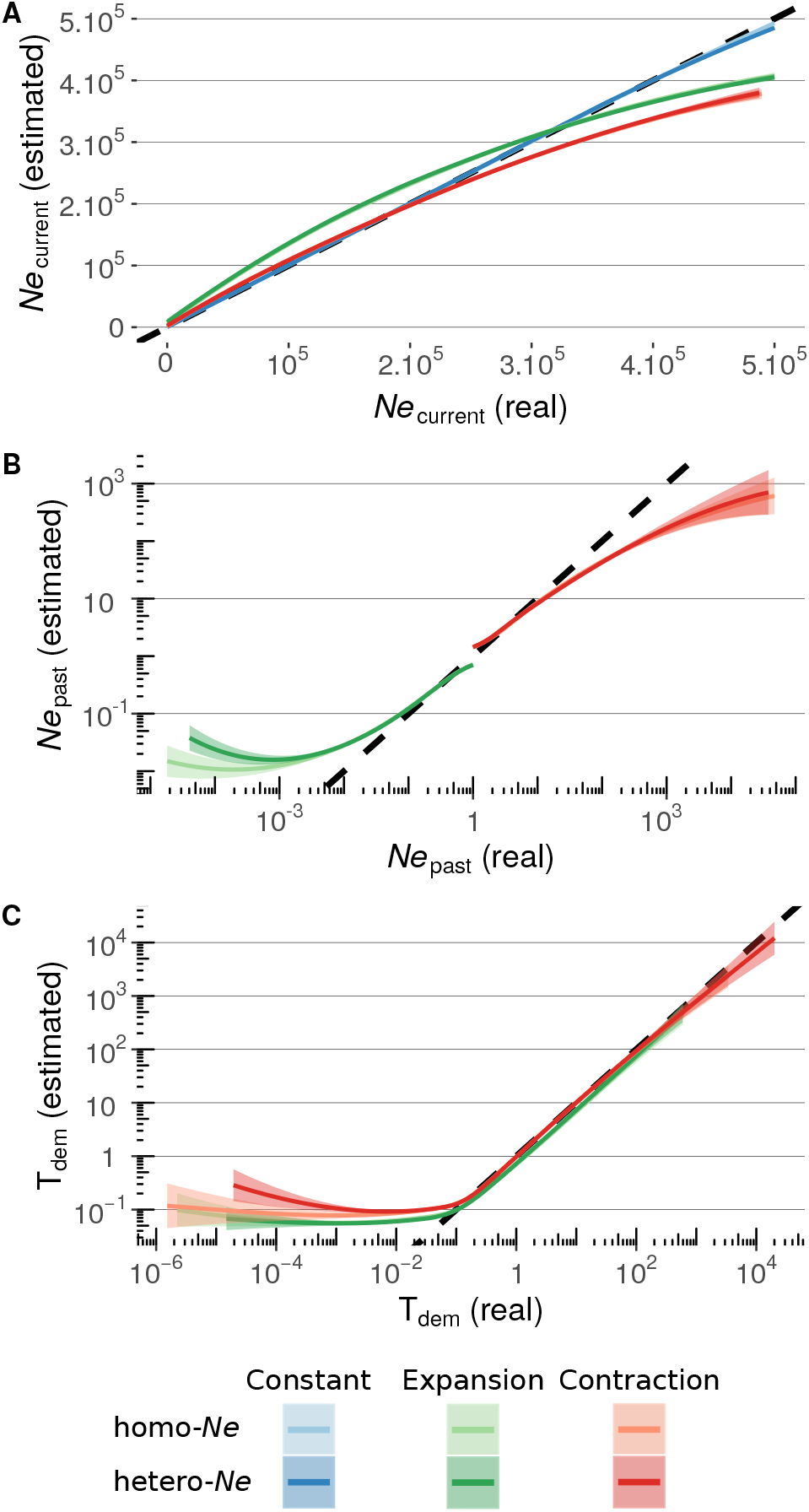
DILS performance for parameter estimations of single-population models. 10, 000 pseudo-observed datasets are simulated by taking random parameter values (x-axis) under the six models. These parameters are estimated using DILS (y-axis). The lines represent the loess (locally estimated scatterplot smoothing) regressions between exact and estimated parameter values for each of the six models. The fields represent the 99% confidence interval and the dotted line represents *x* = *y*. Estimation of the effective size of the current population *Ne*_current_ (**A**), of the ancestral population *Ne*_past_ (**B**) and the time of demographic changes *T*_dem_ (**C**). *Ne*_past_ and *T*_dem_ are both expressed here in terms of *Ne*_current_ individuals.

In a similar manner, the quality of inferences of the age of demographic change *T*_dem_ is highly dependent on its relative value with *Ne*_current_ (figure 4-C where the age of the demographic events is expressed as the ratio *T*_dem_/*Ne*_current_). Any change more recent than 0.1*Ne*_current_ generations ago will be dated with poor precision. Conversely, the age of events older than 0.1*Ne*_current_ appears more accurately recaptured by our ABC approach.

#### Two-population models

The error rate in the estimation of the parameter *Ne*_current_ is of the same order of magnitude as in models with a single population (figure 5-A; table S3). However, the imprecision increases with ongoing migration and tends to underestimate *Ne*_current_. Estimates of *Ne*_current_ are thus more accurate for the SI model, than for the AM model, and the worst for IM and/or SC. This negative effect of ongoing migration on the accuracy of parameter estimation is more pronounced for the ancestral population size *Ne*_past_ (figure 5-B). Hence, the ongoing migration implemented in the IM and SC models will lead to the overestimation of very low *Ne*_past_ values and underestimation of large *Ne*_past_ values.

**Fig. 5.**
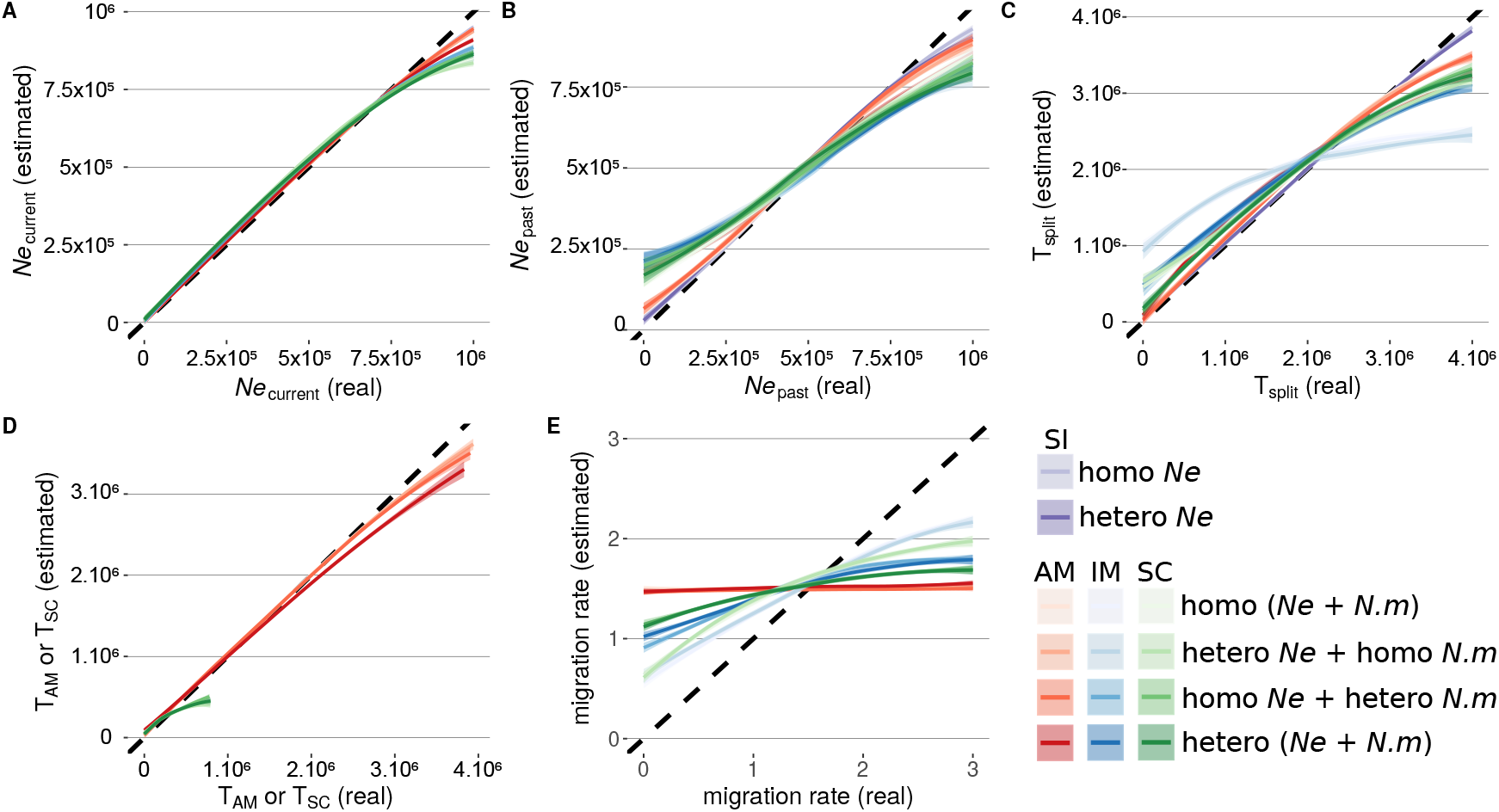
DILS performance for parameter estimations of two-population models. 2,000 pseudo-observed datasets are simulated by taking random parameter values under the 14 models and analyzed using the same procedure as to produce the figure 4, but for the effective size of the current population *Ne*_current_ (**A**), the ancestral population size *Ne*_past_ (**B**), the time of split *T*_split_ (**C**, in generations), the times of demographic transitions *T*_AM_ and *T*_SC_ (**D**) and the migration rate *N.m* (**E**).

The precision of the estimate of *T*_split_ for a given model is of the same order of magnitude as for the ancestral size, with the exception of an accentuated imprecision of *T*_split_ in the IM model when the migration is homogeneous along the genome (figure 5-C; table S3). The AM and SC models both have an additional parameter describing the time of the demographic transition between two periods (with and without migration). In the AM model, *T*_AM_ describes the number of generations during which the two current populations remain genetically isolated after a period of ancestral migration. Conversely, in the SC model, *T*_SC_ describes the number of generations where the two current populations are connected by gene flow during a secondary contact occurring after a past period of isolation. For the AM model, *T*_AM_ is better estimated than *T*_split_ unlike *T*_SC_ under the SC model (figure 5-D; table S3).

Finally, DILS performance to estimate the migration rate *N.m* is reported on the figure 5-E. The poor estimation accuracy for *N.m* contrasts sharply with the reliable inferences obtained when comparing the ‘ongoing migration’ *versus* ‘current isolation’ supermodels (paragraph 3). Indeed, it is straightforward to discriminate between these two categories of supermodels while an accurate estimate of the migration rate is more challenging to obtain (figure 5-E). We were unable to reach a reliable measure of *N.m* for the AM model, but more accurate inferences are obtained for both the IM and SC models. Accuracy is reported to increase for models where *N.m* is homogeneous (table S3). We also investigated how the number of sampled loci can impact the quality of inferences. The results presented above correspond to simulated datasets of 100 loci. When reproduced with 50 loci (tables S4–S6), we show that DILS is not robust enough to study single population models. On the other hand, a reduction in the number of loci does not change the detection of migration in two-population models (although SI versus AM and IM versus SC comparisons are less robust). Errors associated with parameter estimates are of the same order of magnitude for 50 as for 100 loci.

### Detection of barriers to gene flow

One additional purpose of DILS is to identify genomic regions which are associated with a local reduction in the introgression rate *N.m*. This analysis will only be carried out by DILS if the observed dataset is better explained by 1) a demographic model with ongoing migration (IM or SC) and 2) a genomic sub-model with gene flow heterogeneity (hetero-*N.m*). To achieve this purpose, DILS will infer the parameters under the model that best explains the data. Then, for each locus, DILS performs a comparison between two models that differ only for the parameter *N.m*: 1) the *migration* model corresponds to the whole set of parameters estimated under the best supported model; 2) the *isolation* model corresponds to the previous model whose *N.m* has been set to zero, because a barrier gene impedes gene flow locally along the chromosome. Therefore, such a locus should be supported by the isolation model if the barrier effect is strong. This approach therefore seeks to approximate a continuous variable, *N.m*, by a dichotomous choice of model: region with a local-isolation *versus* local-migration.

In order to evaluate DILS performance, the locus-specific model comparison was applied for locus simulated under an IM model with different values of *N.m* in [0, 1] (figure 6). A value of zero means no exchange during the divergence process from one population to another. A value of 1 means that there is one immigrant individual on average every generation. We simulated loci 10, 000 times for different combinations of *N.m* and *T*_split_ under an IM model. Then, for each simulated dataset, we applied the locus-specific model comparison to finally record for each locus which model is the best between local-migration and local-isolation. Ideally, we aim that DILS considers 100% of the simulations with *N.m* = 0 as local-isolation, and 100% of the simulations with *N.m* = 1 as local-migration. Values of *N.m* greater than 1 were not explored because the comparison between “high migration” and “very high migration” is not relevant here.

**Fig. 6.**
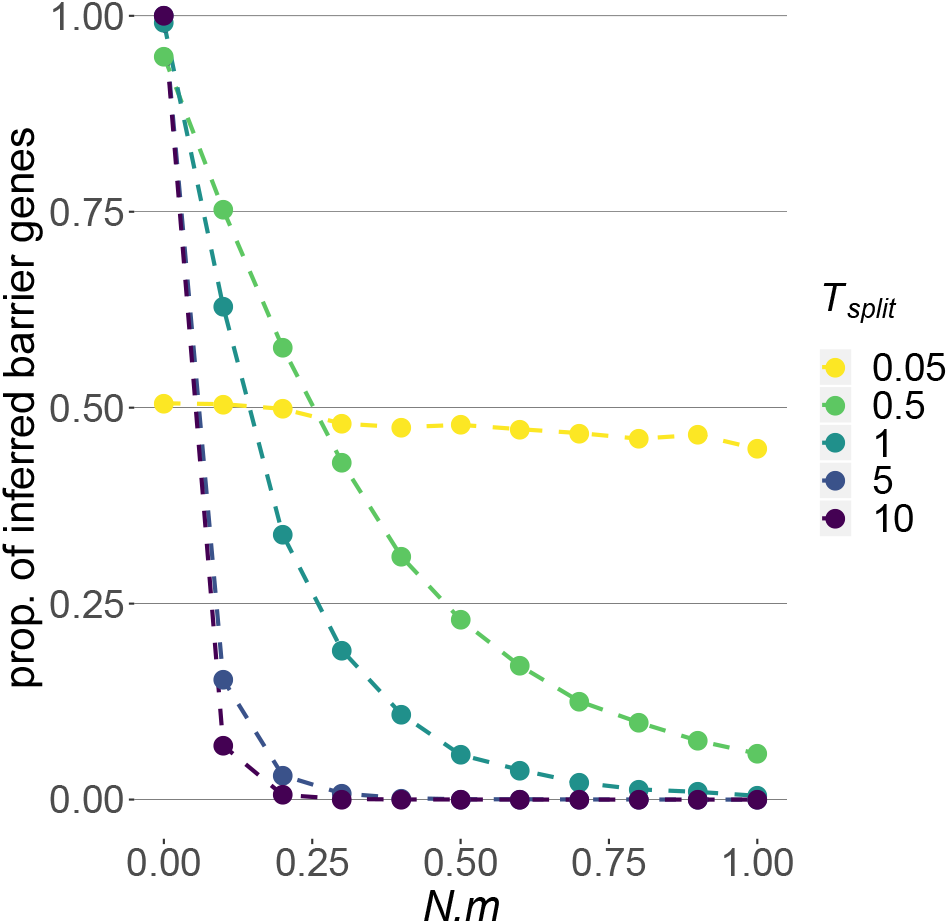
Detection of barriers to gene flow. x-axis: 11 explored values of the locus-specific *N.m* migration rate under an IM model. y-axis: proportion of loci supported by DILS as being linked to a barrier to gene flow (i.e. for which the best model corresponds to local-isolation in the locus-specific model comparison). The colors designate five different divergence times of the IM model (*T*_split_, figure 1). The unit time is in *Ne* generations where *Ne* is the number of haploid individuals making up the population. If *Ne* = 100, 000 individuals, then *T*_split_ = 5 means a divergence time of 500, 000 generations under the IM model. If *Ne* is the number of diploids, then *T*_split_ must be multiplied by two to find the same relationship. Each combination of *T*_split_ and *N.m* was independently simulated 10, 000 times and analyzed by DILS to get the proportion of loci best fitting the local-isolation model for a given combination of parameters. The estimated points are connected by dotted lines for visibility.

As shown in figure 6, in the case of two populations of 100, 000 individuals separated only 5, 000 generations ago (*T*_split_ = 0.05), DILS will support gene flow for ≈ 50% of the loci that have a *N.m* equals to zero (figure 6). For *N.m* = 1, the proportion of loci inferred as local-isolation is of similar magnitude, indicating that DILS is not at all designed to detect barriers to gene flow in the genomes of populations that have separated very recently. Our recommendation, therefore, is to disregard the results of DILS if the studied populations are extremely recent. However, as soon as barrier regions have enough time to differentiate (*T*_split_ ≥ 0.5; figure 6), then ≈ 100% of loci with *N.m* = 0 are correctly inferred as local-isolation, and only few loci with *N.m* = 1 are incorrectly supported by the model of local-isolation. DILS performance therefore depends directly on the true history of the studied populations/species, not on the amount of data. The ideal case for identifying which loci in the genome are linked to barriers occurs when the patterns of polymorphism and divergence at such loci differ greatly from the rest of the genomic background (figure 6). An ideal demographic scenario for identifying barriers with DILS would be:

1. a divergence that is old enough to allow the neutral regions linked to barriers to have at least one position with variants that are differentially fixed between the two populations/species.
2. a migration rate in the genomic background high enough to counteract the effect of differentiation in non-barrier regions.

### Illustration of DILS with RNA-seq data from28 pairs of *Mytilus* populations

We now illustrate DILS potential to contribute to the study of speciation among 8 populations of a complex of four *Mytilus* mussels species (figure 7-A; (Bierne et al. 2003; Popovic et al. 2019)). Using DILS, we established the relationship between molecular divergence and genetic isolation over 28 pairs of *Mytilus* populations. The aim was to identify which pairs of populations, characterized by different levels of molecular divergence (net synonymous divergence between 0.003% and 6.705%), are inferred to be currently connected by ongoing gene flow (figure 7-B). This large scale analysis within the same genus was made possible by the use of a large RNA-seq dataset recently published by (Popovic et al. 2019) in 44 individuals of *Mytilus* from 8 populations (one *M. edulis*, five *M. galloprovincialis*, one *M. planulatus* and one *M. trossulus*; figure 7-A).

**Fig. 7.**
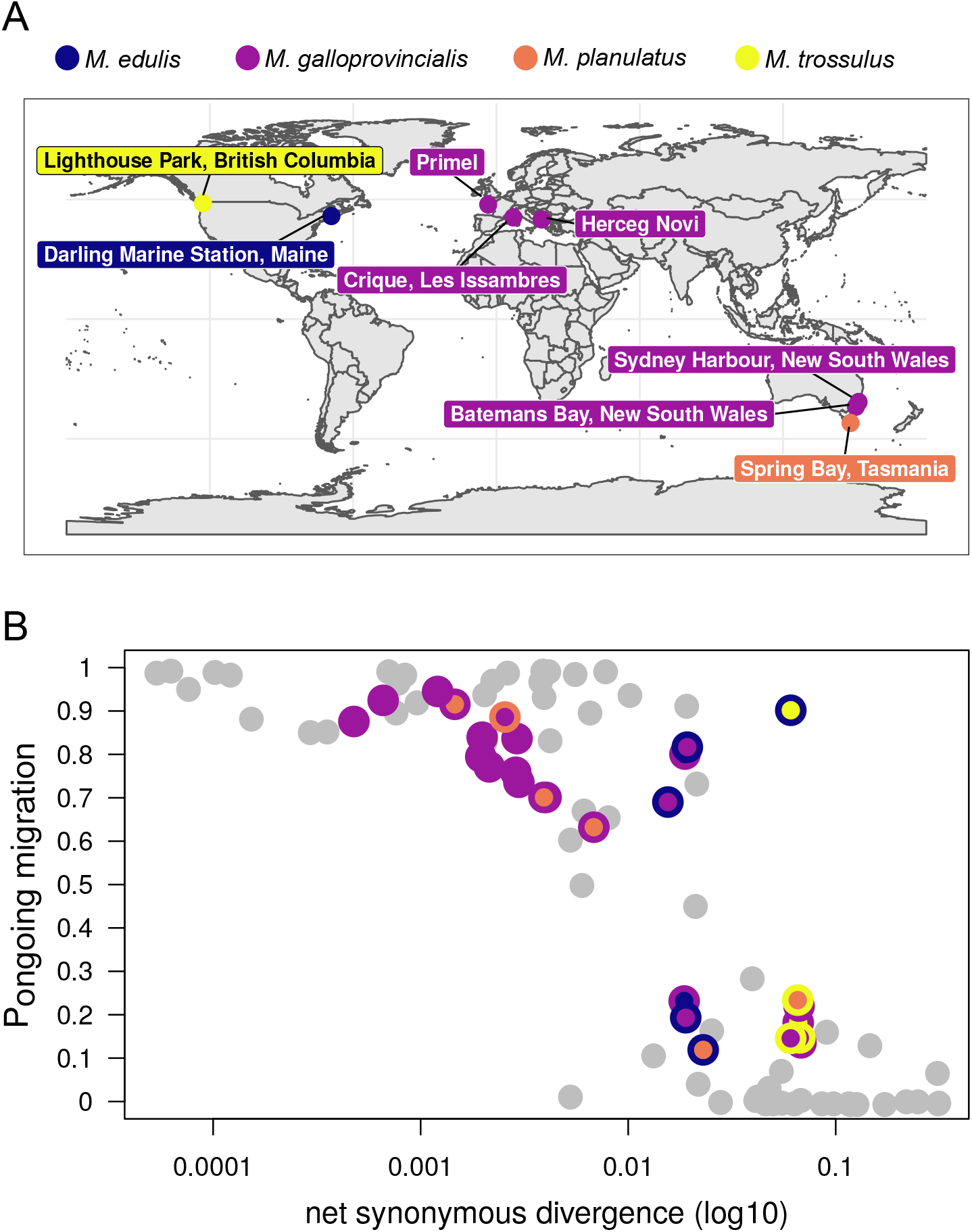
Application of DILS to a *Mytilus* RNA-Seq dataset. **A**: Transcriptomes were obtained in (Popovic et al. 2019) from 44 individuals sampled from 4 labelled species (throughout 8 localities), providing 28 possible pairs of *Mytilus* populations analysed to test for a genetic connection through migration events (SRP218536; https://cutt.ly/ErrDuwj). A median number of 1, 407 coding genes was used to perform demographic inferences after filtering the data (*min* = 144 genes; *max* = 2, 899 genes; depending on the pair of *Mytilus* considered). **B**: The x-axis shows the net divergence measured at the synonymous positions of the sequenced genes. The y-axis shows the probability provided by DILS of models with ongoing migration (IM + SC). Grey dots correspond to the 61 pairs of populations/semi-isolated species/species studied in (Roux et al. 2016). The coloured dots correspond to the 28 pairs of newly analyzed *Mytilus*. The colours refer to the labelled species from which the partners of each designated pair originate.

Out of 28 pairs of *Mytilus* populations that have been tested for ongoing gene flow, 9 pairs receive support for models with current isolation, while models with migration suggest a better fit in the remaining 19 pairs (figure 7-B). Within the group composed of *M. galloprovincialis* and *M. planulatus*, the 15 possible pairs are characterized by levels of divergence ranging from ≈ 0.003% (Crique - Herceq) to ≈ 0.673% (Primel - Spring). All of them are supported by models with ongoing gene flow. Our ABC analysis provides support for gene flow for a single inter-specific pair with high divergence level ≈ 6% (Darling - Lighthouse). This pair is the only one to be genetically connected by migration among the 8 pairs in our analysis that have a level of net synonymous divergence higher than 2%, most probably due to gene flow between West Atlantic *trossulus* and *edulis* populations. Such gene flow also contrasts with an analysis realized for 61 pairs of animal species that had been studied without *a priori* on their speciation history, and for which a threshold of ≈ 2% had emerged above which all inter-specific pairs were currently isolated (figure 7-B; (Roux et al. 2016)).

## DISCUSSION

Our statistical analysis platform, DILS, goes beyond simple summary statistics by explicitly testing evolutionary scenarios in a model-based inference framework. In this sense, it is important for users to be mindful that all biological interpretations must be made in light of the models discussed here. This approach is especially time-wise in speciation genomics as comparative studies of closely-related species are accumulating (e.g. in butterflies: (Cong et al. 2019; Edelman et al. 2019; Martin et al. 2019; Ebdon et al. 2020); in birds: (Peñalba et al. 2019); in fishes: (Malinsky et al. 2018); in plants: (Stankowski et al. 2019)); and so there is a strong demand for efficient and powerful inference tools. Applied to genomic data, DILS will first identify the best demographic model to test for changes in effective size and migration rate over time, then it will identify the best genomic sub-model to test for genome-wide heterogeneity in these parameters (and thus linked selection), and finally it will identify loci most associated with genomic regions locked to gene flow.

### Performances of DILS

For single-population models, DILS is highly efficient at distinguishing the three demographic models (Expansion *versus* Constant *versus* Contraction). It fairly discriminates among the two genomic sub-models (homo-*Ne versus* hetero-*Ne*) for the Expansion and Constant models, but has too much ambiguity to distinguish them in a Contraction model. Current population sizes (*Ne*_current_) are accurately estimated under all three demographic models, as well as the time of size change (*T*_dem_) provided that it is not too recent. However, the past population size (*Ne*_past_) is increasingly overestimated in an expansion model (respectively, underestimated in a contraction model) when the contrast with *Ne*_current_ is increasingly sharp.

We also found that in two-population models, DILS very accurately discriminates between the two supermodels classically tested in speciation (“current isolation” *versus* “ongoing migration”), and it discriminates reasonably well among models with ongoing migration (IM *versus* SC), but quite poorly among models without (SI *versus* AM). Within each demographic model, the two *Ne*-genomic sub-models are fairly discernible; and the same is true for the two *N.m*-genomic sub-models (homo-*N.m versus* hetero-*N.m*) in scenarios of ongoing migration. Parameters are reasonably well estimated in all models (i.e. population sizes *Ne*_current_ and *Ne*_past_, and times *T*_dem_, *T*_AM_ and *T*_SC_); except for the migration rate (*N.m*). The latter is poorly estimated in ongoing migration models, and cannot be evaluated at all when migration happened only in the past (AM).

Therefore it is critical for users to be aware of the limits of DILS; especially when it does accurately discriminate among models and estimate parameter values, and when it does not. For example, the best scenarios for identifying barriers to gene flow is when the genetic signal for these loci strongly contrasts with the rest of the genome, *i.e.* when speciation time is long enough to build-up divergence at barrier regions, and migration rate is high enough to homogenize the genomic background between species. In general, DILS fails to make accurate inferences when divergence or changes in population size have occurred very recently.

Another caveat is that (linked) selection causes demographic parameters to be mis-estimated. This includes the impact of both positive selection (e.g. (Schrider et al. 2016)) and background selection (e.g. (Ewing and Jensen 2016); (Johri et al. 2020)). DILS cannot fully account for this inference bias because of the hierarchical manner in which models are inferred (i.e. the genomic models that test for linked selection are analyzed after demography is inferred).

### Collaborative research

DILS was designed with the objective to facilitate collaborative research in speciation. One major question in the field is to understand how fast reproductive isolation builds-up with divergence between lineages, and so how fast introgression decreases along a continuum of molecular divergence. This relationship has been investigated in 61 pairs of animals (Roux et al. 2016) only providing a partial picture. Here, we extended this work by analyzing genomic data of 28 species/populations of *Mytilus* mussels. Within this specific clade, we found a pattern of non-linear decrease of migration probability with the neutral molecular divergence, similar to what was observed in Roux et al. (2016). However, we also documented ongoing migration between two highly divergent mussel species, hence pushing the grey zone of speciation threshold beyond 2% of net synonymous divergence, maybe due to the outstanding life history traits of mussels (i.e. broadcast spawning, high-dispersal larvae, large effective population sizes and living in a highly connected marine environment).

DILS offers the possibility for the users to participate in this enterprise, and record where their biological model falls within this global speciation picture. Such a global picture of transition from gene flow to no gene flow is necessary for the central problem of species delineation (Hey and Pinho 2012). Although a universal criterion for delineating species seems impossible, as exemplified by the mussel dataset, the idea of defining a grey zone by taxonomic system is promising (Galtier 2019). Thus, our collaborative approach option included in DILS will allow in the future to establish a relationship between molecular divergence and genetic isolation for different taxa, *i.e.* vertebrates, terrestrial plants, algae, etc …, and thus will provide delimitation rules by system.

### Non-detailed features

The raw data can be easily visualized with DILS as a site frequency spectrum and summary statistics across loci. DILS produces comprehensive results for each inference step: 1) the global model comparison to estimate the best demo-genomic model; 2) the locus-specific model comparison to identify barrier loci; and 3) the estimation of parameter values for the best model. To help users interpreting these results, DILS produces a series of goodness-of-fit tests to the data. These tests are performed by simulating under the best model each population genetic statistic calculated in section “Summary statistics” (genomic mean and variance of *π*, *θ*, *F*_ST_, etc.), as well as for each bin of the SFS (or jSFS for two-population models). In addition to an individual test for each summary statistic, a test is also performed from statistics transformed by a PCA following (Cornuet et al. 2008; Cornuet et al. 2014). DILS also provides values for each locus of: 1) each summary statistic; 2) the approximated recombination rate calculated based on the four-gamete rule ((Hudson and Kaplan 1985); implemented in a C-code from (Galtier et al. 2017)); and 3) the posterior probability of being genetically linked to a barrier to gene flow (for two-population models only). These results are outputted as interactive graphics in the application.

Our method is implemented in a user-friendly platform allowing the configuration of the ABC analysis via a graphical interface, its execution and visualization of the results. Detailed information on how to use DILS is provided in the manual. The released version of DILS is currently hosted by the CC LBBE/PRABI, but to ensure full reproducibility and portability on any server, DILS is also packaged in a singularity container freely available at https://github.com/popgenomics/DILS_web. The complete analysis of a dataset (model comparison + parameter estimations + locus-specific tests + goodness-of-fit tests) on the host server takes 2h30 on average.

### Prospects for the future

With the improvement of computational methods, it is now possible to simulate entire chromosomes under the full ancestral process of coalescence and recombination (Kelleher et al. 2016). Combining this type of coalescent simulators with haplotype-based statistics in our ABC framework would be very promising to improve estimates of the timing and extent of gene-flow after secondary contact (Harris and Nielsen 2013). The architecture of DILS has been designed to easily add simulators other than *ms* and its modified versions (Hudson 2002). Thus, it would be readily achievable to use forward-in-time simulations including direct selection (Haller and Messer 2019), and information on local recombination rates or gene density, and therefore making inferences for any selective scheme while taking into account the demographic history of the sample, without changing the pipeline upstream or downstream of the simulator. Recent progress in this direction has been made by Johri et al. (Johri et al. 2020) who jointly inferred past demography and background selection using an ABC framework, and by Sackman et al. ((Sackman et al. 2019)) who developed a similar approach to estimate site-specific selection coefficients under a multiple-merger coalescent model.

## DATA ACCESSIBILITY

The web platform for DILS is available at http://eep.univ-lille.fr/en/productions/dils-software. Source codes to deploy DILS can be freely used from GitHub: https://github.com/popgenomics/DILS_web. A manual is also provided. This GitHub repository also contain the *Mytilus* sequences analyzed (sub-directory ‘example’). The corresponding raw data are available under the accession number SRP218536 (https://cutt.ly/ErrDuwj).

## ACKNOWLEDGEMENTS

We are largely in debt to ‘Pôle Rhône-Alpes de Bioinformatique’ who kindly offered to host DILS, in particular Bruno Spataro and Stephane Delmotte. We would also like to acknowledge Cynthia Riginos for the approach to make public a large amount of produced data, and the associated metadata facilitating their processing. This project benefited greatly from discussions with Arthur Weyna and Pierre Barry, and the redaction from LATEX tips by Julien Fraïsse. CF was supported by an Austrian Science Foundation grant (Project M 2463-B29).

## SUPPLEMENTARY MATERIAL

**Supplementary Table 1.**
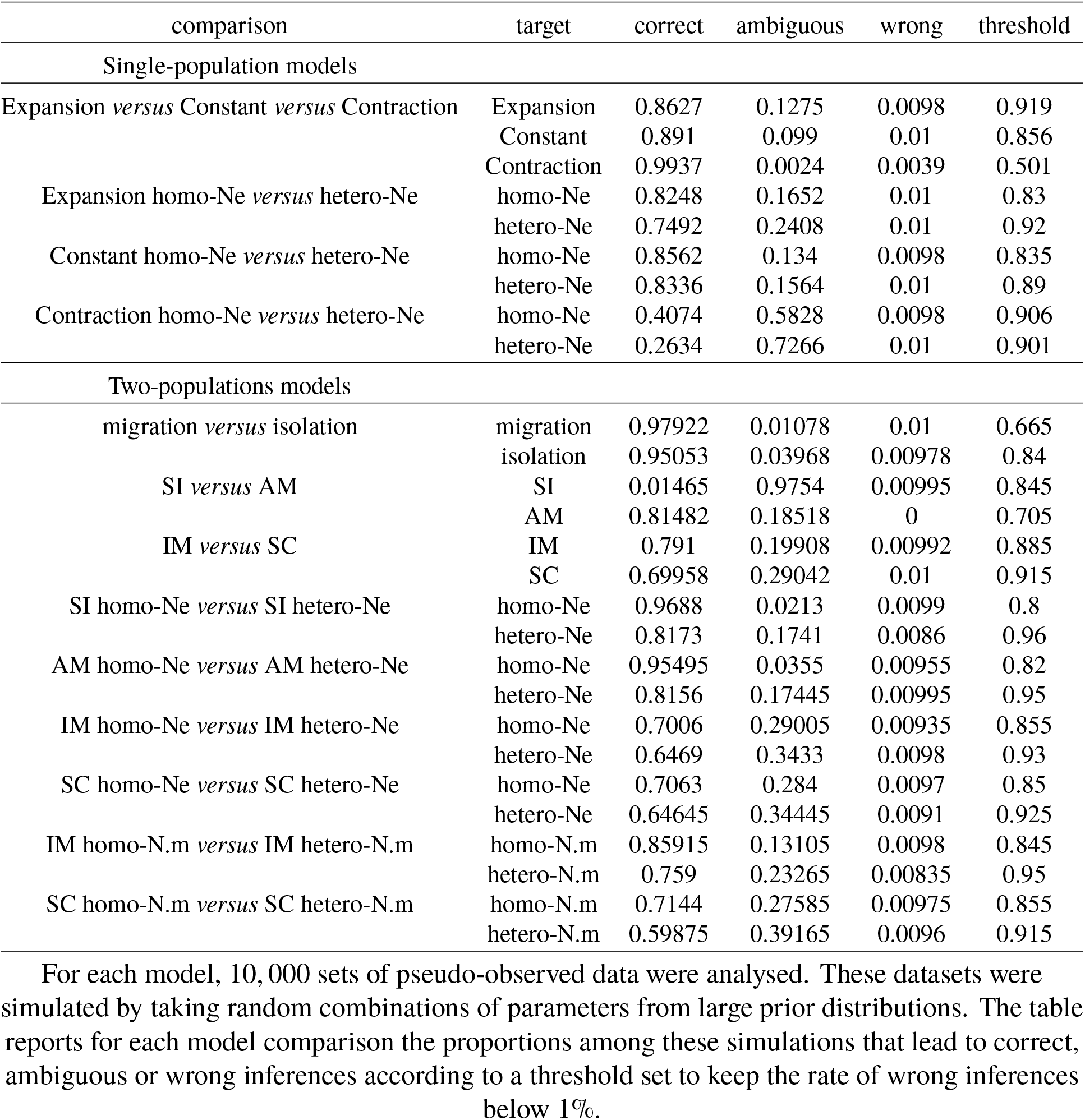
DILS performance for model comparisons

## SUPPLEMENTARY MATERIAL

**Supplementary Table 2.** Mean-squared error in parameter estimations for single-population models

| demographic model | genomic model $Ne$ | $Ne_{\text{current}}$ | $Ne_{\text{past}}$ | $\alpha$ | $\beta$ | $T_{\text{dem}}$ |
| --- | --- | --- | --- | --- | --- | --- |
| Constant | homo | 0.00523 |  |  |  |  |
|  | hetero | 0.01178 |  | 0.39888 | 0.92462 |  |
| Expansion | homo | 0.2009 | 0.00922 |  |  | 0.74168 |
|  | hetero | 0.20754 | 0.01369 | 0.46283 | 0.94578 | 0.74924 |
| Contraction | homo | 0.12258 | 0.08716 |  |  | 0.88229 |
|  | hetero | 0.12563 | 0.09423 | 0.52025 | 0.95338 | 0.90631 |

Estimation errors are calculated on $N = 10,000$ simulated datasets for each model, by using random combination of parameter values. The reported values are measured as follows

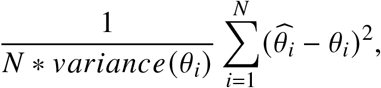

where 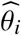 *θ*_*i*_ represent the estimated and the true parameter values respectively

**Supplementary Table 3.**
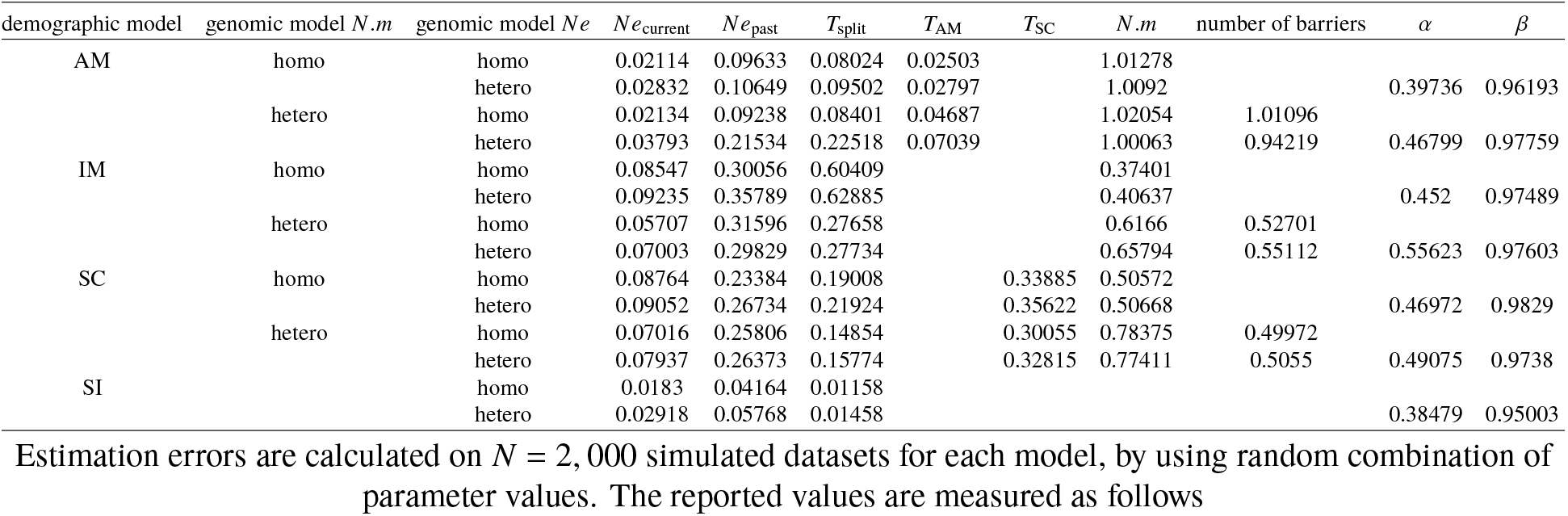
Mean-squared error in parameter estimations for two-population models

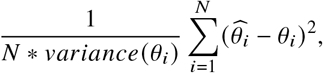

where 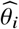 *θ*_*i*_ represent the estimated and the true parameter values respectively

**Supplementary Table 4.** DILS performance for model comparisons (50 loci)

| comparison | target | correct | ambiguous | wrong | threshold |
| --- | --- | --- | --- | --- | --- |
| Single-population models |  |  |  |  |  |
| Expansion *versus* Constant *versus* Contraction | Expansion | 0.3631 | 0.627 | 0.0099 | 0.872 |
|  | Constant | 0.1156 | 0.8744 | 0.01 | 0.924 |
|  | Contraction | 0.8014 | 0.1887 | 0.0099 | 0.801 |
| Expansion homo-Ne *versus* hetero-Ne | homo-Ne | 0.438 | 0.552 | 0.01 | 0.8 |
|  | hetero-Ne | 0.259 | 0.731 | 0.01 | 0.922 |
| Constant homo-Ne *versus* hetero-Ne | homo-Ne | 0.1938 | 0.7962 | 0.01 | 0.898 |
|  | hetero-Ne | 0.3798 | 0.6106 | 0.0096 | 0.896 |
| Contraction homo-Ne *versus* hetero-Ne | homo-Ne | 0.0388 | 0.9512 | 0.01 | 0.847 |
|  | hetero-Ne | 0.1422 | 0.8478 | 0.01 | 0.798 |
| Two-populations models |  |  |  |  |  |
| migration *versus* isolation | migration | 0.96594 | 0.02419 | 0.00988 | 0.83 |
|  | isolation | 0.86258 | 0.12775 | 0.00967 | 0.94 |
| SI *versus* AM | SI | 0.01675 | 0.974 | 0.00925 | 0.835 |
|  | AM | 0.74541 | 0.25459 | 0 | 0.7 |
| IM *versus* SC | IM | 0.14212 | 0.84825 | 0.00962 | 0.815 |
|  | SC | 0.33162 | 0.65912 | 0.00925 | 0.84 |
| SI homo-Ne *versus* SI hetero-Ne | homo-Ne | 0.8025 | 0.1878 | 0.0097 | 0.865 |
|  | hetero-Ne | 0.7669 | 0.2234 | 0.0097 | 0.945 |
| AM homo-Ne *versus* AM hetero-Ne | homo-Ne | 0.8567 | 0.1339 | 0.0094 | 0.845 |
|  | hetero-Ne | 0.76395 | 0.22755 | 0.0085 | 0.945 |
| IM homo-Ne *versus* IM hetero-Ne | homo-Ne | 0.6174 | 0.3733 | 0.0093 | 0.85 |
|  | hetero-Ne | 0.6078 | 0.38245 | 0.00975 | 0.925 |
| SC homo-Ne *versus* SC hetero-Ne | homo-Ne | 0.5762 | 0.41425 | 0.00955 | 0.86 |
|  | hetero-Ne | 0.6026 | 0.38815 | 0.00925 | 0.92 |
| IM homo-N.m *versus* IM hetero-N.m | homo-N.m | 0.82115 | 0.16975 | 0.0091 | 0.84 |
|  | hetero-N.m | 0.7286 | 0.2616 | 0.0098 | 0.94 |
| SC homo-N.m *versus* SC hetero-N.m | homo-N.m | 0.6874 | 0.3026 | 0.01 | 0.84 |
|  | hetero-N.m | 0.5733 | 0.41685 | 0.00985 | 0.9 |

For each model, 10,000 sets of pseudo-observed data were analysed. These datasets were simulated by taking random combinations of parameters from large prior distributions. The table reports for each model comparison the proportions among these simulations that lead to correct, ambiguous or wrong inferences according to a threshold set to keep the rate of wrong inferences below 1%.

**Supplementary Table 5.** Mean-squared error in parameter estimations for single-population models (50 loci)

| demographic model | genomic model $Ne$ | $Ne_{\text{current}}$ | $Ne_{\text{past}}$ | $\alpha$ | $\beta$ | $T_{\text{dem}}$ |
| --- | --- | --- | --- | --- | --- | --- |
| Constant | homo | 0.0228 |  |  |  |  |
|  | hetero | 0.03449 |  | 0.70947 | 0.98946 |  |
| Expansion | homo | 0.08809 | 0.52517 |  |  | 0.16218 |
|  | hetero | 0.35216 | 0.04191 | 0.81278 | 0.99991 | 0.88967 |
| Contraction | homo | 0.47073 | 0.21153 |  |  | 0.62715 |
|  | hetero | 0.46802 | 0.23797 | 0.93637 | 1.02559 | 0.61804 |

Estimation errors are calculated on $N = 10,000$ simulated datasets for each model, by using random combination of parameter values. The reported values are measured as follows

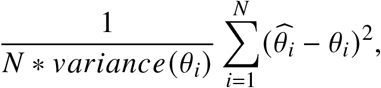

where 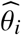 *θ*_*i*_ represent the estimated and the true parameter values respectively

**Supplementary Table 6.**
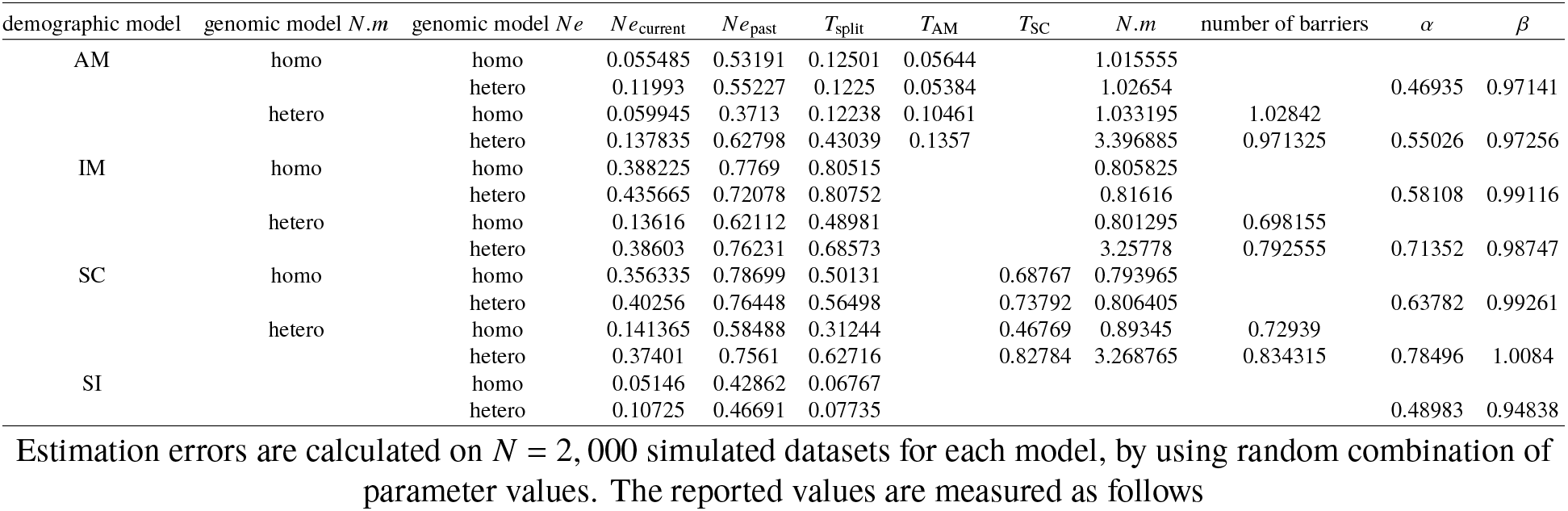
Mean-squared error in parameter estimations for two-population models (50 loci)

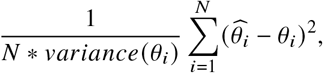

where 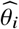 *θ*_*i*_ represent the estimated and the true parameter values respectively

## List of Figures

1 Demographic models currently implemented in DILS

**A.** Single-population models. Demographic changes occurring *T*_dem_ generations ago are modeled backwards in time by sudden transitions from *Ne*_current_ to *Ne*_past_, either for expansions or contractions.

**B.** Two-population models. The Strict Isolation (SI) and Ancient Migration (AM) models are characterized by an absence of ongoing migration. Conversely, the Isolation with Migration (IM) and Secondary Contact (SC) models describe two populations that are currently connected by introgression events at rate *N.m*. The two-population models shown here are of constant size, but DILS optionally incorporates alternative versions of the same four models where effective size can change independently in both daughter populations between the present time and *T*_split_. This is a relevant addition given the influence of over-time size-changes on demographic inferences in speciation scenarios (Momigliano et al. 2020). . . . . . 43

2 DILS performance for hierarchical comparison of single-population models

DILS first performs a comparison of the three demographic models (Expansion *versus* Constant *versus* Contraction). In a second step, it compares two genomic sub-models (homogeneous *versus* heterogeneous genomic distribution of *Ne*) for the best supported demographic model. The pie charts designate for each model the proportion of simulations performed under the corresponding model that is strongly and correctly captured (correct: blue), strongly and incorrectly captured (wrong: yellow) and without strong statistical support for any of the studied models (ambiguous: purple). The performance of DILS was based on 10, 000 pseudo-observed datasets for each of the Expansion / Constant / Contraction demographic models. Each of these 10,000 simulated datasets are evenly distributed between the two genomic sub-models, homo and hetero *Ne*. The parameters used for the simulated datasets are randomly drawn from uniform laws, with *Ne* in [1-1,000,000] individuals and *T*_dem_ in [1-2,000,000] generations. Each simulated dataset consists of 100 loci of length 1, 000 nucleotides. . . . . . . . . . . . . . . . . . . . . . . . 44

3 DILS performance for hierarchical comparison of two-population models

Two-population analyses are performed in three steps (panel **A**). DILS first performs a comparison of the two supermodels (“Current isolation” versus “Ongoing migration”). In a second step, it compares two demographic models for the best supported supermodel: if ‘current isolation’, then DILS tests SI versus AM; if ‘ongoing migration’, then DILS tests IM versus SC. In a third step, DILS compares genomic sub-models for variation of effective population size, *Ne* (panel **B**) and migration rate, *N.m* (panel **C**; only for ongoing migration models) thus testing for linked selection. The letters ‘o’ and ‘e’ in panels B and C indicate simulations performed under genomic h**o**mogeneity and h**e**terogeneity models, respectively. The pie charts designate for each model the proportion of simulations performed under the corresponding model that is strongly and correctly captured (correct: blue), strongly and incorrectly captured (wrong: yellow) and without strong statistical support for any of the studied models (ambiguous: purple). The performance of DILS was based on 10, 000 pseudo-observed datasets for each of the SI / AM / IM / SC demographic models. Each of these 10,000 simulated datasets are evenly distributed between the four genomic sub-models, homo and hetero *Ne* or *N.m*. The parameters used for the simulated datasets are randomly drawn from uniform laws, with *Ne* in [1-1,000,000] individuals, *N.m* in [0-20] 4*.N.m* units where *m* is the fraction of each population made up of new migrants each generation, *T*_split_ in [1-2,000,000] generations and *T*_dem_ / *T*_SC_ / *T*_AM_ between *T*_split_min_ and the sampled *T*_split_ value. Each simulated dataset consists of 100 loci of length 1, 000 nucleotides. . . . . . . . . . . . . . . . . . . . . . . . . . . . . . . . . . . . . . . . . . . . . 45

4 DILS performance for parameter estimations of single-population models

10, 000 pseudo-observed datasets are simulated by taking random parameter values (x-axis) under the six models. These parameters are estimated using DILS (y-axis). The lines represent the loess (locally estimated scatterplot smoothing) regressions between exact and estimated parameter values for each of the six models. The fields represent the 99% confidence interval and the dotted line represents *x* = *y*. Estimation of the effective size of the current population *Ne*_current_ (**A**), of the ancestral population *Ne*_past_ (**B**) and the time of demographic changes *T*_dem_ (**C**). *Ne*_past_ and *T*_dem_ are both expressed here in terms of *Ne*_current_ individuals. . . . . . 46

5 DILS performance for parameter estimations of two-population models

2, 000 pseudo-observed datasets are simulated by taking random parameter values under the 14 models and analyzed using the same procedure as to produce the figure 4, but for the effective size of the current population *Ne*_current_ (**A**), the ancestral population size *Ne*_past_ (**B**), the time of split *T*_split_ (**C**, in generations), the times of demographic transitions *T*_AM_ and *T*_SC_ (**D**) and the migration rate *N.m* (**E**). . . . . . 47

6 Detection of barriers to gene flow

x-axis: 11 explored values of the locus-specific *N.m* migration rate under an IM model. y-axis: proportion of loci supported by DILS as being linked to a barrier to gene flow (i.e. for which the best model corresponds to local-isolation in the locus-specific model comparison). The colors designate five different divergence times of the IM model (*T*_split_, figure 1). The unit time is in *Ne* generations where *Ne* is the number of haploid individuals making up the population. If *Ne* = 100, 000 individuals, then *T*_split_ = 5 means a divergence time of 500, 000 generations under the IM model. If *Ne* is the number of diploids, then *T*_split_ must be multiplied by two to find the same relationship. Each combination of *T*_split_ and *N.m* was independently simulated 10, 000 times and analyzed by DILS to get the proportion of loci best fitting the local-isolation model for a given combination of parameters. The estimated points are connected by dotted lines for visibility. . . . . . . . . . . . . . . . . . . . . . . . . . . . . . . . 48

7 Application of DILS to a *Mytilus* RNA-Seq dataset

**A**: Transcriptomes were obtained in (Popovic et al. 2019) from 44 individuals sampled from 4 labelled species (throughout 8 localities), providing 28 possible pairs of *Mytilus* populations analysed to test for a genetic connection through migration events (SRP218536; https://cutt.ly/ErrDuwj). A median number of 1, 407 coding genes was used to perform demographic inferences after filtering *the data (*min* = 144 genes; *max* = 2, 899 genes; depending on the pair of Mytilus* considered). 49

**B**: The x-axis shows the net divergence measured at the synonymous positions of the sequenced genes. The y-axis shows the probability provided by DILS of models with ongoing migration (IM + SC). Grey dots correspond to the 61 pairs of populations/semi-isolated species/species studied in (Roux et al. 2016). The coloured dots correspond to the 28 pairs of newly analyzed *Mytilus*. The colours refer to the labelled species from which the partners of each designated pair originate.

